# A Custom Microcontrolled and Wireless-Operated Chamber for Auditory Fear Conditioning

**DOI:** 10.1101/738336

**Authors:** Paulo Aparecido Amaral Júnior, Flávio Afonso Gonçalves Mourão, Mariana Chamon Ladeira Amâncio, Hyorrana Pereira Pinto, Vinícius Rezende Carvalho, Leonardo de Oliveira Guarnieri, Hermes Aguiar Magalhães, Márcio Flávio Dutra Moraes

## Abstract

Animal behavioral paradigms, such as classical conditioning and operant conditioning, are an important tool to study the neural basis of cognition and behavior. These paradigms involve manipulating sensory stimuli in a way that learning processes are induced under controlled experimental conditions. However, the majority of the commercially available equipment did not offer flexibility to manipulate stimuli. Therefore, the development of most versatile devices would allow the study of more complex cognitive functions. The purpose of this work is to present a low-cost, customized and wireless-operated chamber for animal behavior conditioning, based on the joint operation of two microcontroller modules: Arduino Due and ESP8266-12E. Our results showed that the auditory stimulation system allows setting the carrier frequency in the range of 1 Hz up to more than 100 kHz and the sound stimulus can be modulated in amplitude, also over a wide range of frequencies. Likewise, foot-shock could be precisely manipulated regarding its amplitude (from ~200 μA to ~1500 μA) and frequency (up to 20 pulses per second). Finally, adult rats exposed to a protocol of cued fear conditioning in our device showed consistent behavioral response and electrophysiological evoked responses in the midbrain auditory pathway. Furthermore, the device developed in the current study represents an open source alternative to develop customized protocols to study fear memory under conditions of varied sensory stimuli.

## INTRODUCTION

Animal behavioral paradigms have long played an important role in understanding the underlying neurobiological mechanisms of learning and memory processes; ubiquitously considered one of the greatest challenges in Neuroscience (Squire, 2009). In Classical or Pavlovian Conditioning, animal innate or reflex responses become evocable by stimuli that are usually neutral (such as sound or visual stimuli), if previously paired with an emotionally relevant stimuli, characterizing the basis of an associative learning process (Kim and Jung, 2006). Using a more appropriate terminology, the conditioned responses (CR) are established under an appropriate contingency of unconditioned stimulus (US) presentations paired with the conditioned stimulus (CS) occurrences (Baron, 1959; Holland, 1977). Proper controls undergo the exact same procedure aside from the fact that CS or US can be presented alone or CS is not paired in time with the US (Rescorla, 1967). Although the laboratory equipments designed to perform such associative learning protocols are supposedly fairly simple, the price can be prohibitive for small budget projects and they are usually quite inflexible in terms of controlling and programming the stimuli. The latter may constitute a drawback for the design of customized behavioral paradigms which may use amplitude modulated stimuli in order to isolate neural circuitry involved in the sensory processing by means of steady state evoked responses (Lockmann et al., 2017; Pinto et al., 2017). In addition, the study of more complex cognitive and behavioral processes becomes impracticable, since they require more sophisticated and robust means of controlling contextual parameters and interacting with the animals (Cushman et al., 2013).

Fortunately, custom development of laboratory tools is becoming more feasible with the advances in technology (Buccino et al., 2018; Pineño, 2014; Siegle et al., 2017; Sinard and Gershkovich, 2012; White et al., 2019). Mainly, low-cost commercially available microcontroller modules have recently achieved a high level of integration and processing power, encouraging their acquisition and use in research laboratories. Arduino, for example, is an attractive hardware and software open-source solution (D’Ausilio, 2012) to program several different families of microcontrollers for a number of different consumer needs (Pineño, 2014; Teikari et al., 2012).

Considering this scenario, the purpose of our work is to present a customized, low-cost, microcontrolled and wireless-operated device for animal behavior conditioning. We propose a scalable architecture, based on Arduino Due board and ESP8266-12E module, which allows adding different sensory stimuli and behavioral feedback aside from what we chose to depict in this work (i.e. programmable sound stimulation waves and electric shock). Likewise, other elements, such as sensors, actuators and touchscreens, can be easily integrated to the apparatus by connecting them to innumerous general-purpose input/output (GPIO) pins available, or through digital communication protocols, like I2C, UART or SPI. This allows the design of a low-cost widely equipped apparatus and eases the development of customized behavioral paradigms.

## METHOD

### SYSTEM OVERVIEW

The main electronic components of the custom conditioning chamber are the ESP8266-12E module (https://www.adafruit.com/product/2491) and the Arduino Due board (https://store.arduino.cc/usa/due). The first one is an internet-of-things device from Espressif company, available since 2014. It has a Tensilica L106 32-bit microcontroller and an 80 MHz CPU clock. It operates at 3.3 V and supports WiFi (IEEE 802.11 g/b/n), I2C, SPI and UART communication protocols. Eleven GPIO pins and 32 kB RAM are available in the module. The ESP8266 is used here mainly as a programmable and flexible user interface, presented on a web page format, to allows users to write and read parameters to the device without the need of physically implementing buttons, knobs and displays.

The Arduino Due is a 32-bit microcontroller board based on the Atmel SAM3X8E ARM Cortex-M3 CPU. It has a 84 MHz CPU clock, 54 digital input/output pins, 12 analog inputs, 2 DAC (digital to analog) and can communicate through SPI, I2C, UART and USB.

Both modules are programmed with an Arduino sketch (*Web_Page.ino* for the ESP8266-12E and *Stimuli.ino* for the Arduino Due) with specific assignments and they cooperate to produce all the apparatus functionality. The modules communicate with each other via a 16-bit serial peripheral interface (SPI) and through digital signaling via two GPIO pins. In the scope of this work, systems for auditory and electrical stimulation (footshock) were implemented, stimuli that can be used as CS and US, respectively, in behavioral paradigms (Figure 1).

**Figure 1.**
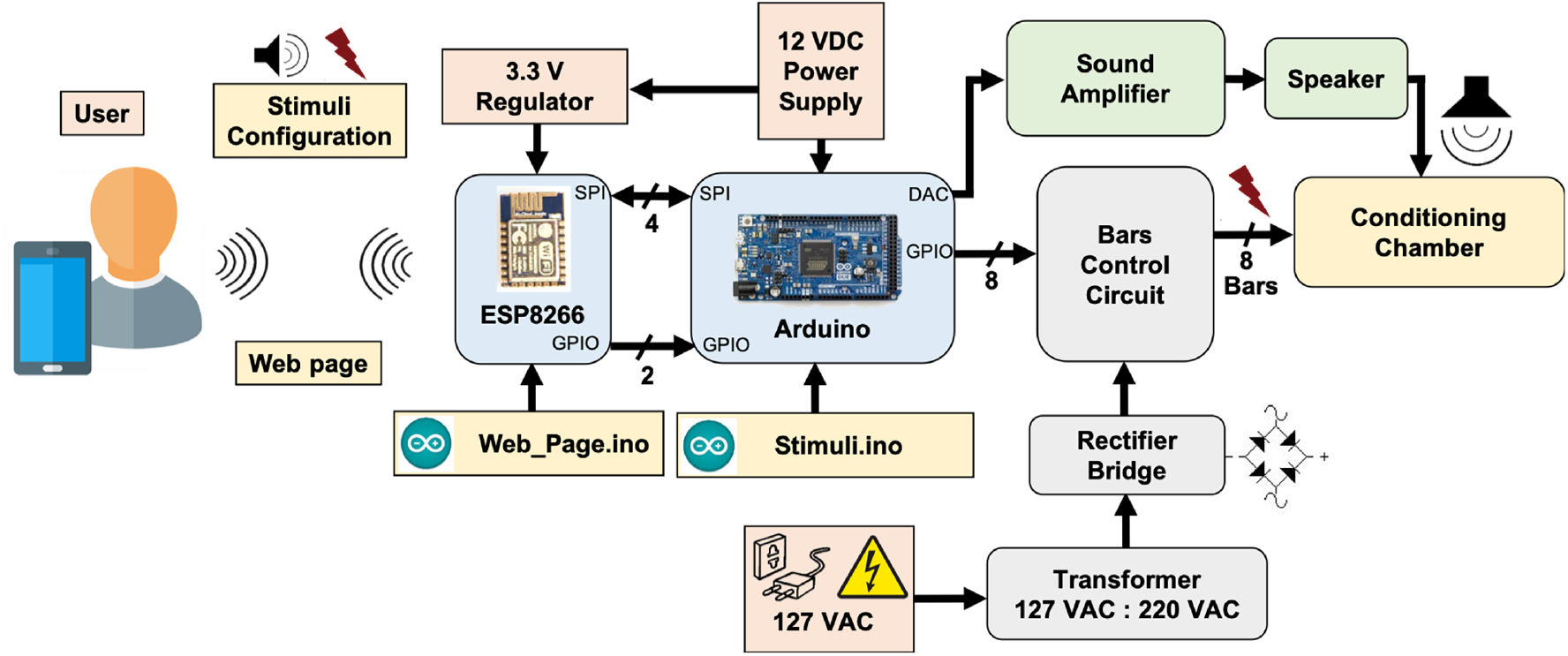
System overview. ESP8266-12E module and Arduino Due board cooperates in order to generate custom modulating tones and aversive stimulation, based on footshocks applied through electrified bars, programmed via user interface.

The auditory stimulating system consists of a 12-bit digital-to-analog converter (DAC; available at Arduino Due board), a commercial audio amplifier and a speaker. The footshock system, in turn, is formed by a high DC-voltage (310 V_DC_) supply circuit in series with a current limiting resistor, significantly higher than the animal’s resistance, in series with a variable resistor in order to deliver a short pulse of constant current through the bars on the chamber. The footshock system was designed to limit stimulus intensity between ~200 μA and ~1500 μA and can be manually set by a potentiometer and an ammeter. Embedded electronics were designed to have only one bar under the animal, at a specific time, serving as current sink. The rate that bars are "scanned" and the duration of each shock pulse are programmable. The user interface for operating the apparatus was developed in a web interface, implemented in HTML within the *Web_Page.ino* sketch, making it not only very flexible and comprehensive in terms of controlling system parameters. Parameters such as sound and shock onset (seconds), shock pulses intervals between bars (milliseconds to seconds), sound intensity (%), sound carrier frequency (Hz) and modulating frequency (Hz) can be remotely programmed to perform the experiments. In addition, each added trial can be saved by simply insert the parameters into a sequence and loaded it in later experimental sessions (Figure 2). Thus, behavioral paradigms can be easily configured using iPads, smartphones, notebooks or any device with a Wi-Fi connection.

**Figure 2.**
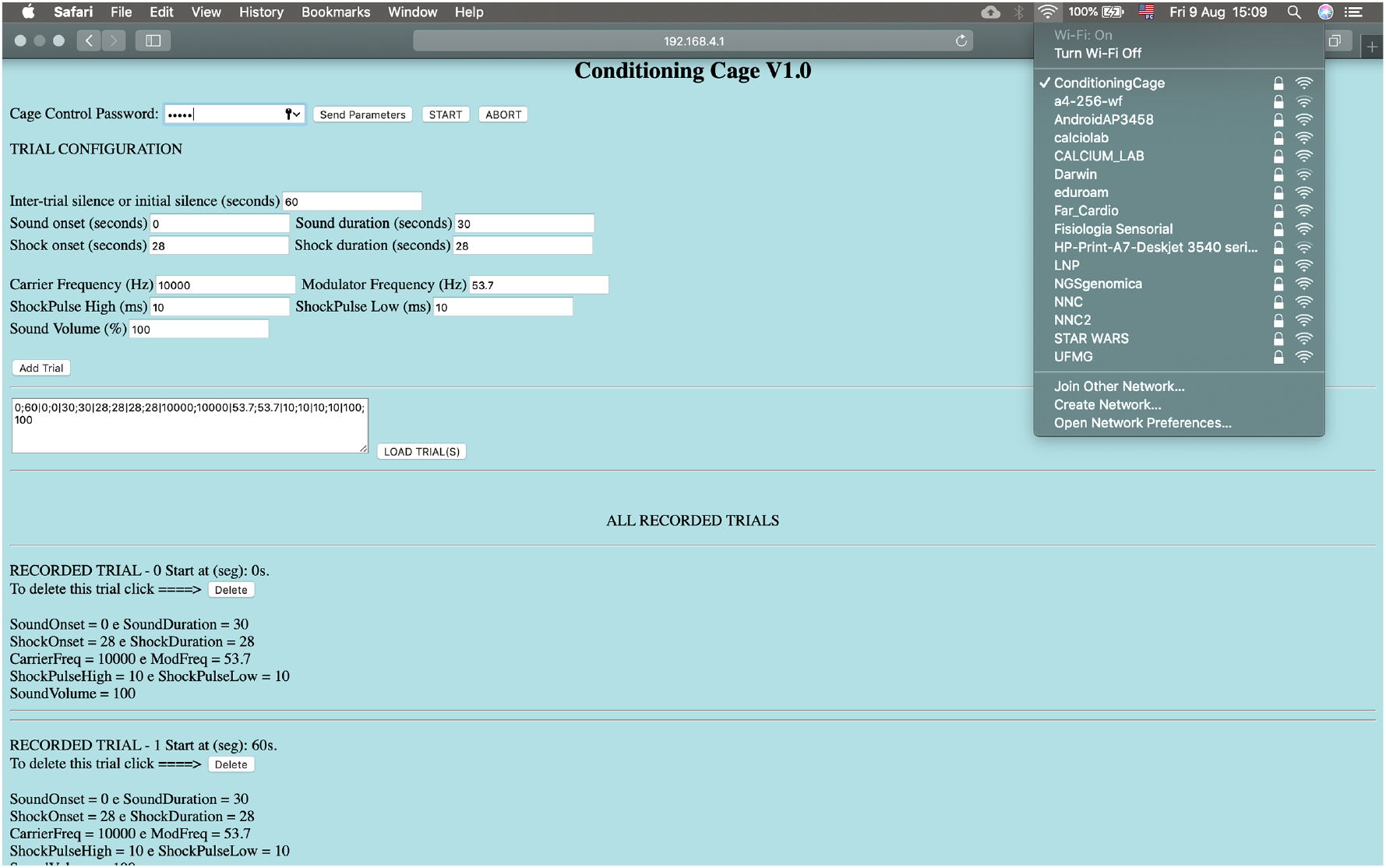
The conditioning chamber can be remotely controlled by a HTML interface. Parameters: Sound and shock onset (seconds), shock pulses intervals between bars (milliseconds to seconds), sound intensity (%), sound carrier frequency and modulating frequency. Each added trial can be saved by simply insert the parameters into a sequence and loaded it in later experimental sessions. Wi-Fi connection: ConditioningCage, IP address: http://192.168.4.1, Password: 0112358132, Cage control password to send the parameters: 12345. Start / Abort: Start and stop experiment.

#### Hardware Design

The custom conditioning chamber dimensions are 330 x 200 x 300 mm^3^ (length, width and height, respectively), built with 5 mm-thick transparent acrylic boards. An acrylic board covers the chamber, preventing the animals from escaping the context (Figure 3A-C). The chamber floor is built with 16 stainless steel cylindrical bars, 250 mm-long and 5 mm in diameter each. They are arranged in parallel, spaced 20 mm from each other and placed through small bilateral holes with 0.6 cm in diameter. The number of bars can be increased up to 32 and spaced 7.5 mm from each other.

**Figure 3.**
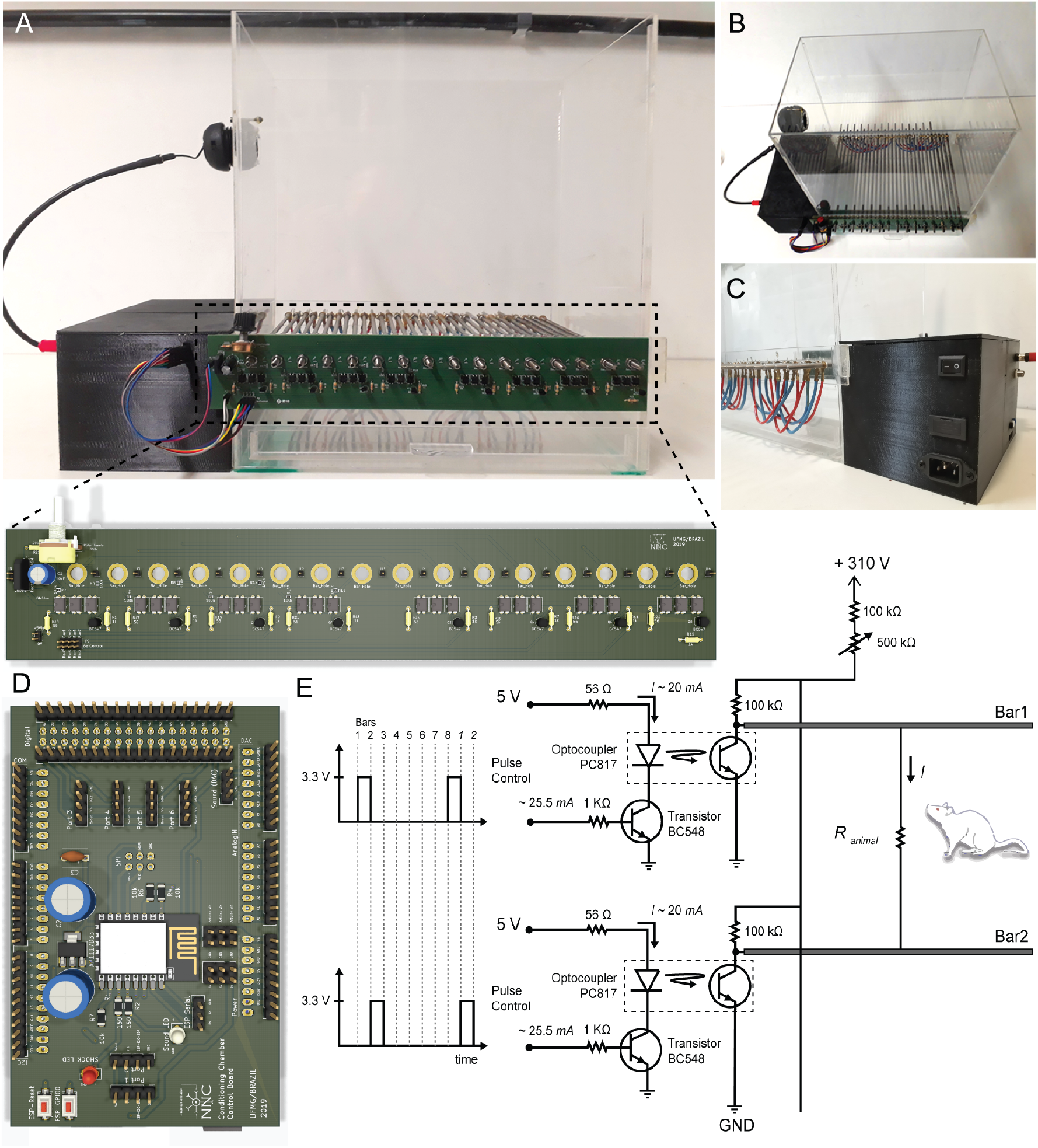
Hardware Design. A, *Top*) Conditioning chamber front view. *Bottom*) Two-layer printed circuit board with power circuit. Attached to the chamber via 16 mounting holes, this board allows creating electric potential between the conductive bars. B and C) Up view and back view of the conditioning chamber respectively. Detail for black box built on the 3d printer where all electronic components are organized. D) Two-layer printed circuit board with control circuit. It has ESP8266-12E footprint, Arduino Due shield and it outputs a low power audio waveform and digital signals with temporal patterns for electric stimulus control. E) Electric potential on each bar is controlled through a switch (npn transistor operating in saturation mode). A set of optocouplers are used in order to isolate power circuit from control circuit. Output voltage is 0 V_DC_ or 310 V_DC_. Output current ranges from ~200 µA to ~1500 µA.

In order to deliver the shock through the bars, a 127 V_AC_: 220 V_AC_ transformer (0.03 kVA) raises and isolates the AC voltage from the electric grid. A diode rectifier bridge (2KBP06M; 600 V / 2A) and electrolytic capacitor (10 μF / 350 V) at the rectifier output produce a constant voltage of approximately 310 V_DC_. The high voltage end passes through a current limiting resistor, set to a maximum of 1-2 mA, and then through a variable resistor in order to "fine tune" the output current. The current remains fairly stable since both resistors are within an order of magnitude higher than the conductance changes expected from the animal stepping on the bars and closing the circuit. Finally, a series of simple common-emitter circuits, using bipolar transistors (BC547B) as switches are used to drive optocouplers (PC817) in order to deliver a current pulse to each individual bar, therefore guaranteeing that whatever combination of bars the animal is stepping on will still deliver the foot shock. The bars are sequentially "scanned" at a programmable interval and the pulse duration can also be set by the user. The Arduino Due board outputs a set of digital signals that control each bar potential. Altogether, this constitutes an aversive stimulating system for US presentations (Figure 3E).

A 12-bit digital-to-analog converter at Arduino Due is used to generate analog waveform. The algorithm behind signal generation is the following: 1) a set of 8 points (2 bytes each) form the template of a sinusoidal waveform - or any period of a programmable signal - that will be repeated at the carrier frequency. Thus, the sampling frequency (fa) of the D/A output will be 8 times that of the carrier frequency (fc). 2) The amplitude of each 8 sample block will be proportionally given by each byte of a sequence of bytes representing the modulation frequency (fm). That is, the sequence of bytes is composed by a minimum of fa/(fm. 8) bytes repeated up to the duration set by the user. In this way, we were able to generate quite a variety of stimulation waveforms at a very low computational cost (within the range of allowing ultrasound stimulation) and using few parameters and/or memory (Figure 4A-D).

**Figure 4.**
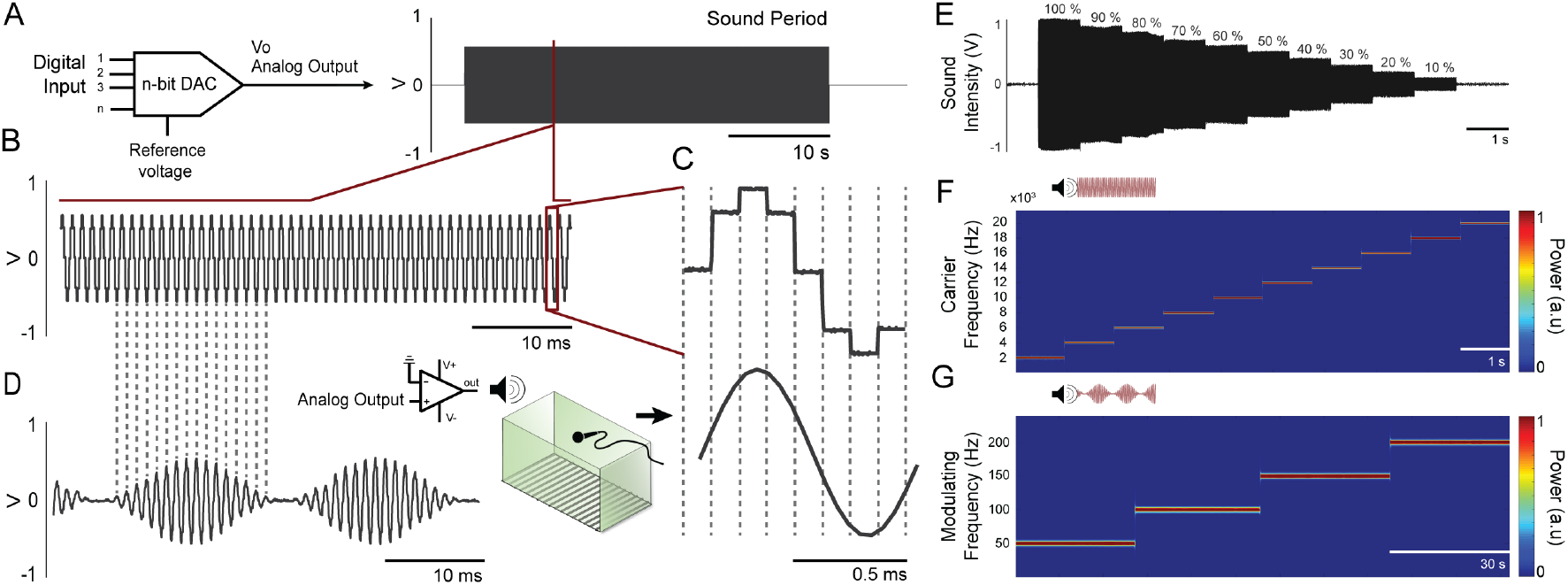
Waveforms can be generated from digital-to-analog converter at Arduino Due. A) A 12-bit digital-to-analog converter (DAC) is used to generate custom audio waveforms. B) Tones with adjustable frequency can be produced with 8 voltage steps within each tone cycle. Although these voltage steps introduce undesirable high frequency components on audio signal, these components can be attenuated at power amplifier circuitry, since its frequency response is similar to a low-pass filter. C) Upper waveform shows a single cycle generated at the DAC output, where the 8 voltage steps are visible. Lower waveform shows the same cycle measured at power amplifier output, which low-pass filter the audio signal and outputs a smooth waveform, practically a pure tone. D) Amplitude modulation can be applied to the audio waveform, with adjustable modulating frequency. E) The amplitude of the programmed tone It can be set to various levels. F) Time-frequency power of the carrier frequency in a range of 2 kHz - 20 kHz (steps by 2 kHz) were calculated by the short-time Fourier transform-STFT demonstrating a constant amplitude over time. G) Time-frequency power of the modulating frequency envelope in a range of 50 Hz - 200 Hz (steps by 50 Hz) were calculated by the short-time Fourier transform-STFT demonstrating a constant amplitude over time.

#### Firmware Design

Arduino software applications were implemented for operating and controlling the custom chamber: *Web_Page.ino*, uploaded to ESP8266-12E module, and *Stimuli.ino*, uploaded to Arduino Due board.

Programmed with *Web_Page.ino* sketch, ESP8266-12E module takes care of basically two things: (1) it provides the user interface, through which the experimental protocols can be programmed (this interface is a web page, available from an independent access point WiFi network managed by the ESP8266-12E), and; (2) it uses 2 GPIO pins to determine when the Arduino Due should turn the conditioned (CS) and unconditioned (US) stimuli on or off, according to the parameters specified via web interface (Figure 2).

Programmed with *Stimuli.ino* sketch, Arduino Due board takes care of generating CS and US with specific temporal patterns, as specified by user. This routine manages triggers and CPU interruptions in order to generate, at the Arduino DAC output, an analog waveform with carrier and modulating frequencies, as well as a specific amplitude. In addition, GPIO pins are used to control each bar potential to produce custom aversive footshock patterns.

### SYSTEM VALIDATION

The validation of the customized apparatus was made in two steps. At first, bench tests were carried out in order to verify the correct functionality of the circuits, safety specifications and firmware for CS and US presentation before using the system in a living organism. In the second stage, experimental animals were submitted to a classical fear conditioning (CFC) task using Auditory Steady State Stimulation (ASSR - 10 kHz carrier frequency modulated at 53.7 Hz) while recording local field potentials in the Inferior Colliculus (IC), the principal midbrain nucleus of the auditory pathway. The IC recordings show stable oscillations at the same frequency of the stimulus amplitude modulating component (i.e. when stimuli are turned-on), reflecting entertainment of neurons responding to sound. This particular steady-state evoked potential (SSEP) neural response is called an ASSR (Kuwada et al., 1986; Picton et al., 2003).

#### Bench tests

In order to assess the flexibility in programming the sound stimulus, as well as the precision of its configurable parameters (mainly carrier frequency and modulating frequency), different sound stimuli reproduced by the audio system were recorded by means of a microphone and placed inside the customized chamber. The sampling frequency employed was 96 kHz and the raw data were qualitatively analyzed with custom-made and built-in MATLAB codes (MATLAB R2016a). The time-frequency power of the carrier and modulating frequency were calculated by the standard built-in spectrogram function (short-time Fourier transform-STFT; non overlapping, 2,048-point Hamming window). In the case of the modulating frequency, the audio signals were preprocessed so that only the envelope of the signal (thus, the modulating signal) was considered. This was accomplished by means of the built-in Matlab Hilbert transform function.

In order to determine the flexibility and precision of the aversive electrical stimulus, a 20 kΩ resistor was connected between two bars of the chamber, simulating the resistance of a rodent that could eventually be touching them (Figure 6A). Similar values of resistance for rats were reported in previous work (Walters and Tullis, 1966). The voltage drop across this resistor was measured with an oscilloscope (GDS-2202A GW INSTEK) for four different programmed shock values (400 μA, 600 μA, 800 μA and 1000 μA) during one second time course (Figure 6B), so that the electric current could be easily determined using Ohm’s Law.

Additionally, another bench test evaluated how much the footshock intensity changes with respect to variability in animal body conductance (which can occur due to humidity and different body masses between animals). For this, a 400 μA footshock was programmed and its intensity was measured in three different cases: R = 15 kΩ, R = 20 kΩ and R = 25 kΩ (Figure 6D-E).

#### Proof of concept: Classical fear conditioning (CFC) and Local field potential (LFP) recordings

Male Rattus norvegicus (Wistar) weighing 270-310 g were supplied by the *Instituto de Ciências Biológicas 2* (BICBIO 2) vivarium, housed under controlled environmental conditions (22 ± 2 ºC), with a 12:12 h light-dark cycle and free access to food and water. All experiments were approved by the Institutional Animal Care and Use Committee at the *Universidade Federal de Minas Gerais* (CEUA-UFMG; protocol no. 112/2014.), and were conducted in accordance with *Conselho Nacional de Controle de Experimentação Animal* (CONCEA) guidelines defined by Arouca Act 11.794 under Brazilian federal law. CEUA directives comply with National Institutes of Health (NIH) guidelines for the care and use of animals in research.

#### Surgery

Animals were anesthetized intraperitoneally with ketamine/xylazine solution (15 mg/kg and 80 mg/kg, respectively), the head surface was shaved and then the animal was positioned in a stereotaxic frame (Stoelting, Wood Dale, IL). After asepsis with alcohol (70%, topical) and povidineiodine solution (7.5%, topical) a local anesthesia with lidocaine clorohydrate-epinephrine [1% (wt/vol), 7 mg/kg] was applied and an incision in the scalp was made to expose the skull.

Monopolar electrodes for recording, made of a twisted pair of stainless-steel (0.005 in.) teflon-coated wires (Model 791400, A-M Systems Inc., Carlsborg, WA, USA) was slowly lowered and positioned at the ventral region of the left IC central nucleus (AP: −9.0 mm referenced from the bregma, ML: −1.4 mm, DV: −4.0 mm) (Paxinos and Watson 2007). The coordinates for electrode positioning were chosen according to the tonotopic organization of the IC since the ventral layers respond better to the high frequencies (Clopton and Winfield, 1973; Malmierca et al., 2008) as used in the CFC protocol. At the end stainless steel screws were implanted on the nasal bones (anterior to the olfactory bulb), one to the left as the reference electrode (0 V) and another one as the ground. Both reference/ground and recording electrodes were soldiers in a RJ-11 connector, which was fixed to the skull with dental acrylic. Animals received a prophylactic treatment with pentantibiotics (Zoetis Fort Dodge; 19 mg/kg) and anti-inflammatory (Banamine, 2.5 mg/kg) to prevent discomfort and infection after surgery and recovery for 7 day.

#### Classical fear conditioning (CFC)

The CS used in the CFC was a 10 kHz pure tone with its amplitude modulated by a sine wave of 53.7 Hz. These specific tone features were chosen based on previous results (Lockmann et al., 2017; Pinto et al., 2017).

The amplitude modulated tone generated by the customized conditioning chamber was amplified by a commercial sound amplifier (AB100, 100 WRMS, 4 Ω, NCA) and reproduced by a speaker (ST304, 40 WRMS, 8 Ω, Selenium Super Tweeter).

Before the behavioral sessions, the intensity of the CS tone was measured and set to 85 dB SPL at the center of the conditioning chamber (Brüel & Kjær type 2238 sound level meter). This sound pressure level ensures an appropriate electrophysiological response considering the anatomical positioning of the recording electrode (Malmierca et al., 2008; Meeren et al., 2001).

CFC was performed on three consecutive days in two different contexts (Figure 3A). On the first day (preconditioning), ten rats were presented with 5 CS stimuli (30 s each apart from each other, with pseudorandom interval not longer than 120 seconds) in the context A (a black acrylic box, 30 x 20 x 25 cm^3^ with one transparent face and 10% alcohol solution scented).

On the second day (conditioning), the behavioral task took place in the context B, which is the custom chamber proposed in this work (with 0.001% acetic Acid solution scented). The rats were randomly assigned to paired (n = 5) or unpaired (n = 5) groups and similarly to the day 1, five CS stimuli were presented, however five US were applied. The US consisted of a 400 μA current applied through metal bars on the floor over 2 s (Mourão et al., 2016). In the paired group, each presentation of CS was temporally paired with US in the last two seconds of the CS presentations. On the other hand, in the unpaired group, US occurred at random times between CS.

On the third day (retention test), rats were presented to an accurate repetition of the preconditioning procedure. Returning to the context A (10% alcohol solution scented), all animals were presented with 5 CS stimuli (30 s apart from each other, with pseudorandom interval not longer than 120 seconds).

The sessions were recorded by a camera set up in front of the box, and the videos were analyzed by an examiner blind to the experimental group (Figure 5A). Freezing behavior was defined as no movements, except those resulting from breathing, that last a minimum of 3 s (within each 3 s time epoch). Results were expressed as the percentage of freezing during each CS trial (30 s - 10 epochs in which a freezing episode either occurred or not)(Blanchard and Blanchard, 1972; Fanselow and Bolles, 1979; Curzon et al., 2009). Periods outside of the CS presentation were not considered for quantification.

**Figure 5.**
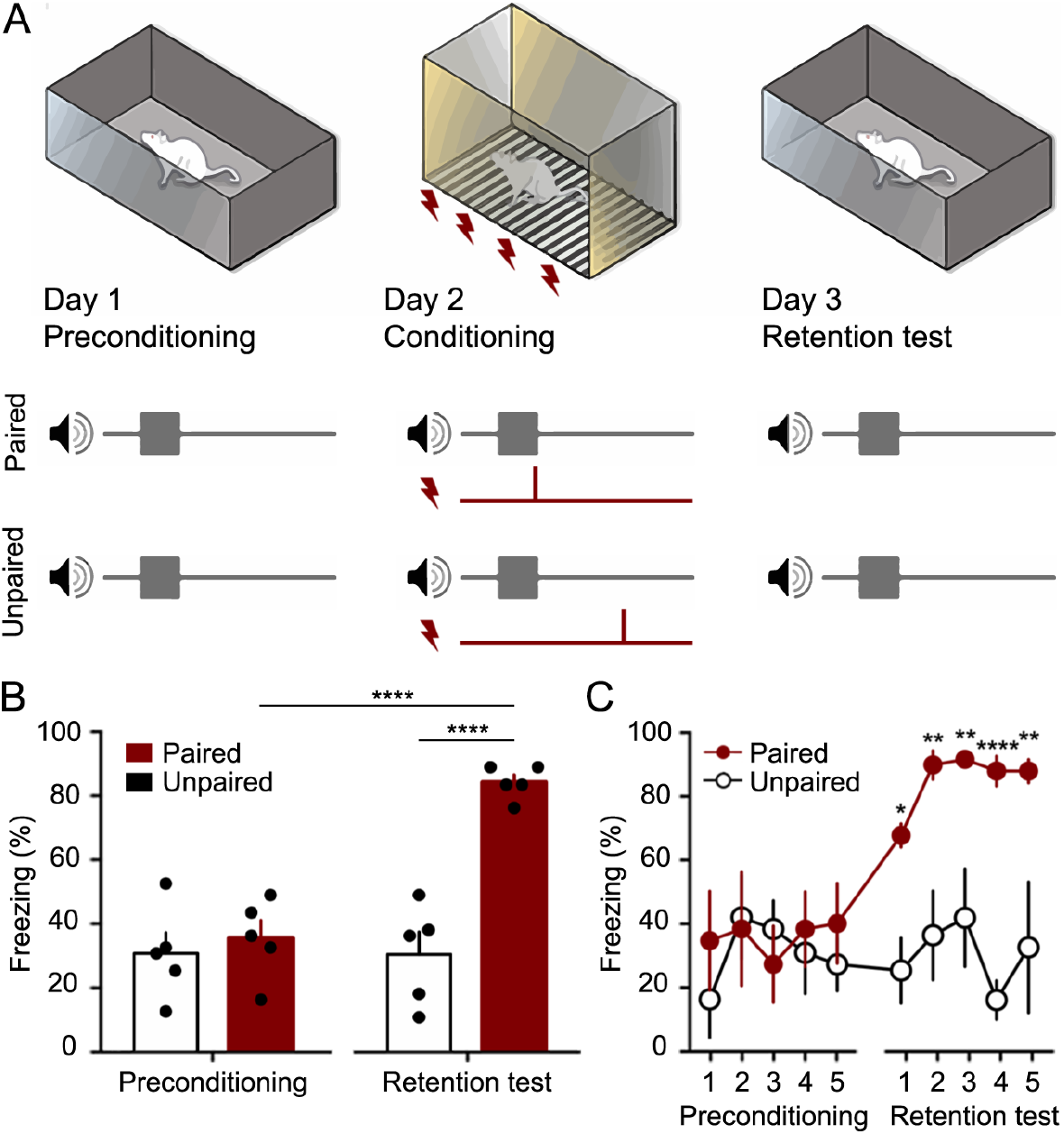
The Fear conditioning protocol (CFC) was performed as a proof of concept for the customized chamber. A) Preconditioning occurred in Day 1, when 5 trials of CS (10 kHz pure tone with its amplitude modulated by a sine wave of 53.7 Hz) were presented to the animals in a context A. The Conditioning occurred in Day 2, when 5 trials of CS and US (a footshock with 400 μA over 2 s) were presented to the Paired and Unpaired groups using the customized chamber. In Day 3, retention test occurred with 5 trials of CS being presented to both groups. Animal fear response (freezing) was measured in order to verify if the association between sound and shock. B) Total mean freezing (± SEM) to CS in preconditioning and test sessions. C) mean freezing (± SEM) to CS over trials. * p < 0.05, ** p < 0.01, **** p < 0.0001. 2-way ANOVA followed by Bonferroni post hoc test.

**Figure 6.**
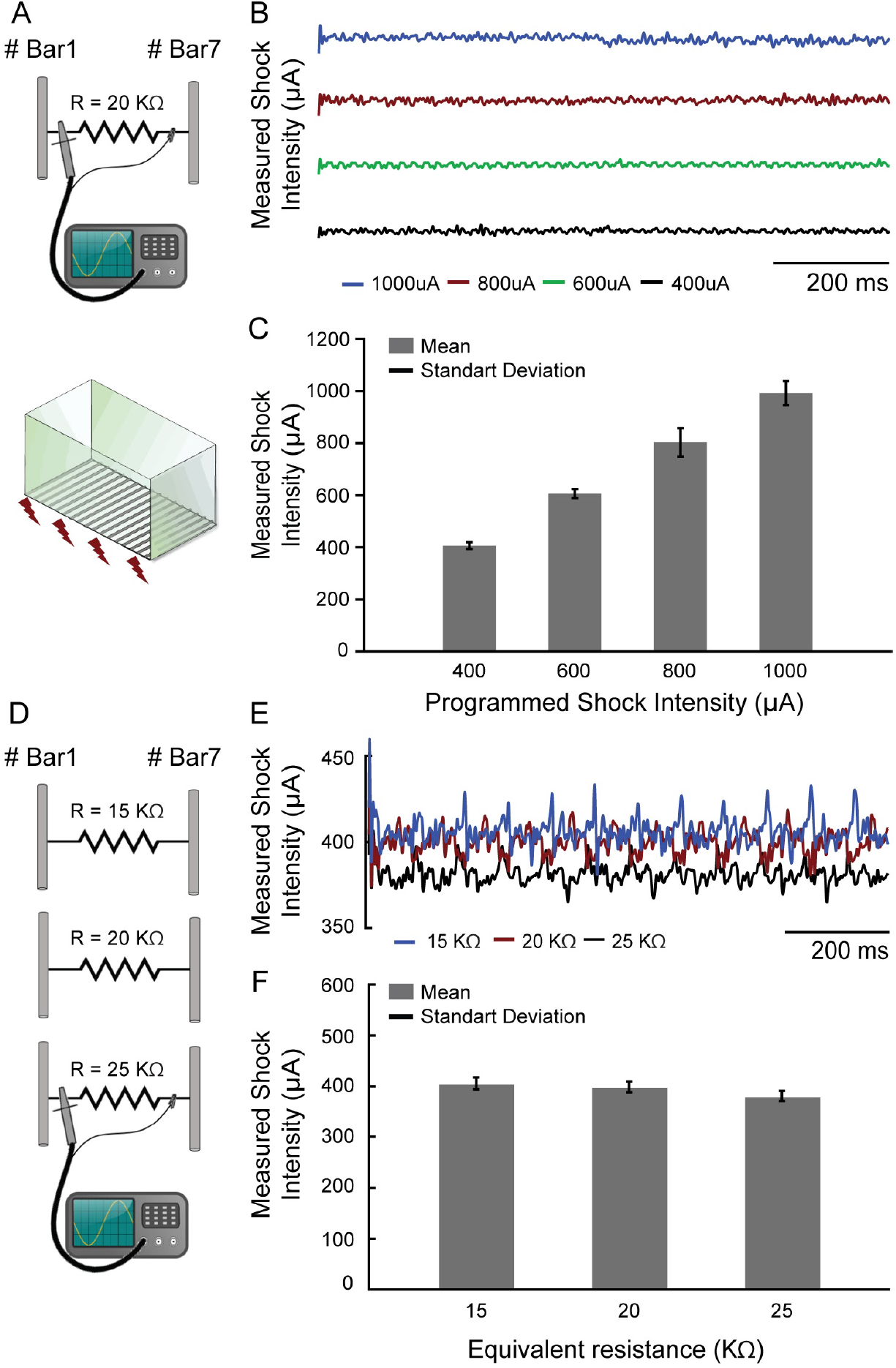
The footshock exhibited precisely configurable patterns. A) The footshock intensity was measured by an oscilloscope (I = V/R) through a resistor that simulates the animal skin resistance. B) The current measurements were made over time in four different programmed shock intensities (μA). C) The mean footshock intensities remaining constant and showed no significant changes. D) The same measurement was performed with three different resistors at a programmed current value to simulate the skin resistance variations. E) The current measurements were made in one programmed value (400 μA) over time. F) The values with a 25 % deviation around the 20 kΩ caused no significant changes in the programmed shock intensities. (maximum ~ 4.74 %, ~ 18 μA, to 25 kΩ).

#### LFP recordings and data analysis

LFP signals from the ventral region of the left IC central nucleus were recorded during the CFC test sessions (Figure 7). The signals were obtained through a pre-amplified unit gain stage (1x gain. ZCA-AMN16 adapter. Omnetics ©. Tucker-Davis Technologies) coupled to a thin recording cable (ZC16 - 16 channel ZIF-CLIP® digital head stages. Tucker-Davis Technologies). A small adapter was built into a printed circuit board so that the commercial pre-amplified unit gain stage coupled to the RJ-11 connector surgically implanted in the experimental animals. The signals were filtered between 1 and 2,000 Hz, amplified by 24,000 V/V and sampled at approximately 12 kHz by a bioamplifier processor (Tucker-Davis Technologies RZ2).

**Figure 7.**
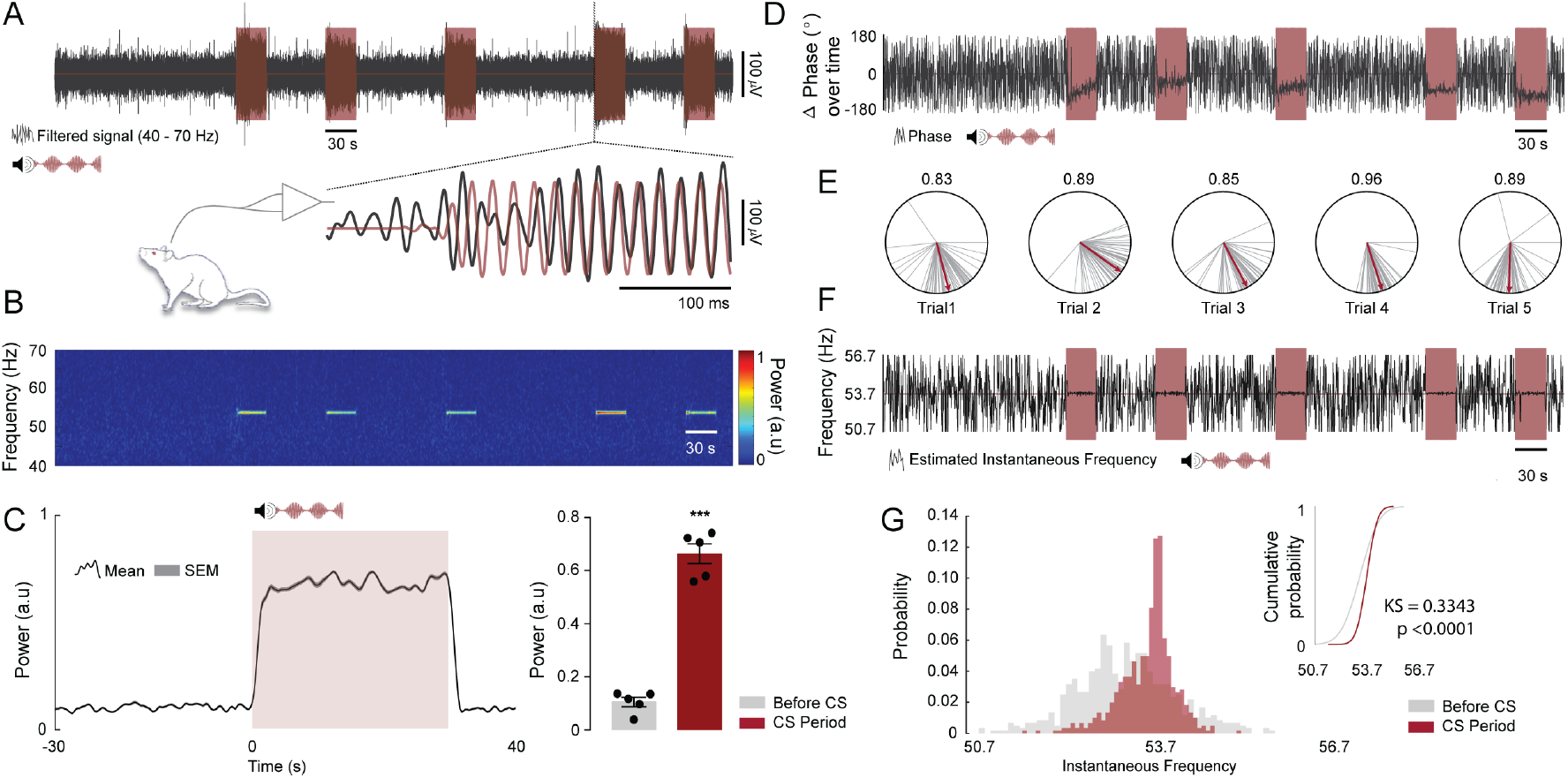
Oscillations in the IC can be entrained by the envelope of an amplitude modulated tone. A, top) Representative time course of LFP (grey) and CS trials (light red, 10 kHz pure tone with an amplitude modulation of 53.7 Hz). A, Bottom) Highlight to a sample time window at the beginning of the fourth trial showing that the IC LFP synchronizes with the CS amplitude envelope. The LFP and the CS envelope were band-pass filtered between 40 and 70 Hz. B) Spectrogram showing SSEP power at 53.7 Hz. (a.u., Arbitrary units). C, left) Mean SSEP power at 53.7 Hz ± 3 Hz over trials for all animals. The figure shows a period of 30 seconds before CS onset, 30 seconds of CS presentation and 10 seconds after. C, right) Qualitative analysis of SSEP power at 53.7 Hz ± 3 Hz for 30 seconds before CS onset compared to the 30 seconds of CS presentation (n = 5 animals; before CS vs. CS period; *t*_(4)_ = 11.42, p = 0.0003). D) Representative time course of delta phase values at 53.7 Hz ± 3 Hz (*ϕSSEP* − *ϕCS*) extracted from the Hilbert coefficients (grey) and CS trials (light red). E) Delta phase vectors computed for 250 ms time windows during CS presentation (grey lines) and the respective estimate mean phase values (red lines). The numbers above the polar plots represents the phase coherence of each trial during CS presentation. F) Representative time course of instantaneous oscillatory frequency estimated at 53.7 Hz ± 3 Hz from the changes in the SSEP phase angles (grey) and CS trials (light red). G) Distribution of instantaneous oscillatory frequencies over trials for 30 seconds before CS onset (light grey) and 30 seconds of CS presentation (light red) and their respective cumulative distributions. Two Sample Kolmogorov–Smirnov test was used to test whether the two samples come from the same distribution (n = 5 animals; before CS vs. CS period delta phase angles distributions; KS = 0.3343, p < 0.0001).

The timestamps locked to the peaks and valleys of the CS modulating frequency were generated in parallel through one of the Arduino Digital I/O pins and then recorded by a digital input port of the bioamplifier processor. Through linear interpolation, these time values were used to obtain an instantaneous phase time series, which in turn were used to reconstruct/estimate the CS modulating signal itself. In this sense the time-frequency analysis could keep engaged with the stimulus presentation.

The data were off-line preprocessed and analyzed with custom-made and built-in MATLAB codes (MATLAB R2016a. EEG lab. *Open source environment for electrophysiological signal processing* - http://sccn.ucsd.edu/eeglab/). Time-frequency power of the SSEP (frequency range 40 - 70 Hz) were calculated by the standard built-in spectrogram function (short-time Fourier transform-STFT; non overlapping, 16,384-point Hamming window).

To calculate the Δ phase between CS amplitude modulating envelope and SSEP, the data were initially filtered at the frequency range of 53.7 ± 3 Hz and the coefficients were extracted by the built-in MATLAB function Hilbert transform. The Δ phase was calculated in the average time windows of 250 ms as the difference between the imaginary components of SSEP and the CS envelope:

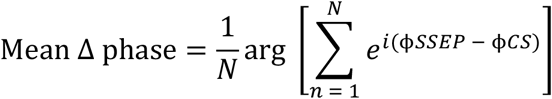

Mean Δ phase: argument of the sum of the phase vectors, where N is the number of time-axis samples of each signal and ϕ*SSEP and* ϕ*CS* are the phase values for the SSEP and CS envelope, respectively.

The phase coherence between CS envelope and SSEP was extracted by means of the length of the average vector of phase angles differences:

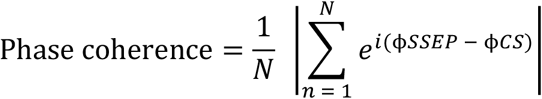

Phase coherence: metric of phase synchronization. Adimensional real value between 0, that represent an uniform phase distribution and 1, a perfect phase grouping (Lachaux et al., 1999; Mormann et al., 2000).

The dynamics of instantaneous oscillatory frequency was estimated from the changes in the SSEP phase angles over time. As described above, the data was filtered (range of 53.7 ± 3 Hz) and the first derivative of SSEP phase from Hilbert coefficients was calculated in the average time windows of 250 ms. The phase-angle time series was then transformed to a time series of instantaneous frequency by multiplying with the sampling rate in hertz and then dividing by 2π (Boashash, 1992; Cohen, 2014).

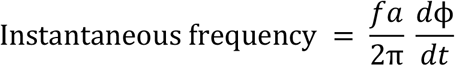

Instantaneous frequency: frequency component linearly proportional present in the modulating signal. Where *fa* indicates the sampling rate and *d*ϕ /*dt* indicates instantaneous variation of unwrapped angles over time.

#### Statistical analysis

The quantitative data are expressed as means ± standard error of the mean (SEM). The approximation to the normal distribution was confirmed by the Kolmogorov– Smirnov test (P > 0.05). Statistical comparisons were made using Student’s t-test or one-way or two-way repeated-measures analysis of variance (ANOVA), followed by Bonferroni post hoc test. For some analyses Two Sample Kolmogorov–Smirnov test was used to test whether two samples come from the same distribution. Values of P < 0.05 were considered statistically significant. Data were analyzed using GraphPad Prism 7.0a Software and MATLAB (2016a), The Mathworks, Natick, USA.

## RESULTS

### Bench test - CS and US stimuli

The auditory stimulation system allows setting the carrier frequency in range of 1 Hz up to more than 100KHz. In addition, the sound stimulus can be modulated in amplitude, also over a wide range of frequencies. Another flexible characteristic of the auditory stimulation is its intensity, which can be programmed by controlling the amplitude of the Arduino digital port output. Some carrier and modulating frequencies were chosen to be depicted in Figure 4. Figure shows that the parameters were adjusted accurately. The amplitude of the programmed tone (in this case 10 kHz carrier frequency) remains stable at its various programmed levels (Figure 4E) and the amplitude of the recorded tones (2 kHz - 20 kHz carrier frequency, Figure 4F; 50 Hz, 100 Hz, 150 Hz and 200 Hz modulating frequency, Figure 4G) remains constant over time.

Regarding the aversive sensory stimulus, it also exhibited precisely configurable patterns, both temporal (number of pulses per second) and intensity (in μA) (Figure 6C). In addition, the footshock system showed little susceptibility to changes on animal body conductance. A 25 % deviation around the 20 kΩ simulated rat resistance caused, in the worst case (R = 25 kΩ), a corresponding variation of only 4.74% (about 18 μA) in the expected shock intensity, which was 400 μA (Figure 6F). We understand that the effect of this on the experimental protocol is negligible and does not harm the behavioral data obtained with the custom box.

It is worth noting that this slight variation in shock intensity was only possible ^because the operating voltage of the shock circuitry is relatively high. The 310 V_DC_^ potential requires resistors of 300 kΩ or more to limit the shock intensity under 1000 μA. Since the animal resistance is much smaller than 300 kΩ, the shock intensity is largely determined by the resistors in the electronic circuitry rather than by the animal itself.

It should be noted that the *Stimuli.ino and Control_Box.ino* sketches can be easily modified to specify which bars will be used to apply footshock. This allows creating different spatial patterns of aversive electrical stimulation, which in turn makes it possible to perform different behavioral tasks.

### Classical fear conditioning with an amplitude modulated tone

Over the past decade, many studies have shown that auditory fear conditioning is able to induce the emergence of defensive behaviors (Maren, 2001). This traditional task can be considered one of the most important methodological tools that allow understanding of the underlying mechanisms of learning and memory (Collins, 2000; Duvarci and Pare, 2014; Maren et al., 2001; Quirk et al., 1995).

In the present work the animals submitted to the CFC in the customized chamber presented a robust conditioned response (Figure 5 B-C). According to the results, in the preconditioning session, the auditory stimulus can be considered a neutral stimulus since it did not elicit high levels of freezing. However, in the test session, 24 h after the conditioning session, the paired group showed a significant increase of Freezing behavior (F_(1,8)_ = 34.26; p < 0.0001), while the unpaired group maintained the baseline values as the preconditioning session. The comparison between the paired and unpaired groups showed significant differences in the test session (F_(1,8)_ = 34.26; p < 0.0001).

### LFP recordings during the CFC Test session

Oscillations in the auditory pathway (Rees et al., 1986) and more specifically in the IC (Lockmann et al., 2017; Pinto et al., 2017) can be entrained by the envelope of an amplitude modulated tone. Furthermore, after an associative learning task in which the auditory stimulus is paired with an aversive stimulus, the temporal dynamics of LFP can change substantially. The phase of evoked oscillation couples to the phase of the amplitude modulated tone and the power significantly increases with the auditory stimulus representations (Lockmann et al., 2017). In this sense, we reproduce the data previously published by our group in an attempt to prove the concept that the developed chamber generates an accurate set of auditory and aversive stimuli.

During the CFC test session, we recorded the oscillatory activity of the ventral region of the IC central nucleus in five rats from paired group. Region that, according to the IC tonotopic organization, perfectly responds to high frequencies (10 kHz - CS auditory stimulus). In addition, we modulate the evoked activity with a specific amplitude envelope (53.7 Hz) to generate a resonant frequency, an SSEP.

As expected, the LFP recorded in the IC shown a distinct power spectral signature embedded in the programmed fm (Figure A-B) and moreover the SSEP power during the mean trials was significantly higher (~3-fold) compared to the previous instants outside the trial periods (Figure 7C. *t*_(4)_ = 11.42; p = 0.0003).

The qualitative analysis of delta phase values extracted from the Hilbert coefficients demonstrates that the angles are organized in clusters without major changes in the mean values. In addition, the data shown a high phase coherence values over each trial, suggesting a high level of synchronization between CS envelope and the SSEP (Figure 7D-E).

Finally, the distribution of instantaneous oscillatory frequencies over trials presented a variation around the modulation frequency, being significantly different from the distribution instantaneous oscillatory frequencies extracted from the previous instants outside the trial periods (Figure 7F-G. KS = 0.3343; p < 0.0001).

## 5. DISCUSSION

The proposed conditioning chamber has an auditory stimulation system which exhibited versatility in programming stimuli characteristics (carrier frequency, modulating frequency, intensity and duration). Furthermore, the equipment also has an aversive stimulation system, based on footshocks applied through electrified bars. Footshock intensity and duration, as well as the number of constant-current pulses per second, can be properly adjusted.

It is important to highlight that the versatility regarding the auditory stimulation system has an important application in providing a characteristic spectral signature in LFP electrophysiological recordings of brain structures processing the sensory stimuli. Lockmann et al (2017) has shown that a pure, continuous and amplitude-modulated tone may acquire biological relevance since the neural activity of inferior colliculus, responsible for one of the first stages of auditory processing, presents significant changes in their temporal dynamics after conditioning. It is important to highlight that the SSEP provides a unique perspective on evaluating circuitry involved in sensory signal processing once its "tags" the electrophysiological recordings of such brain substrates with a distinct spectral signature assigned by the modulation frequency.

In addition, the conditioning chamber architecture allows adding new components to its design with little effort, for example: infrared sensors, touchscreens, servomotors (i.e driving water and food dispensers), among others. This increases the equipment usability as a tool for customized scientific protocol design.

As a high-level programming language platform, and with all community support via forums, the Arduino IDE also enables collaborative effort to enhance and enrich the chamber software routines (*Web_page.ino* and *Stimuli.ino* sketches). Our project is being made available through open-source initiatives https://github.com/fgmourao/NNC_repository) and may be modified by collaborators adding new features to both hardware and software. In this publication all the files and tutorials will be available as supplementary materials.

The initiative to build custom equipment encourages the development of unique investigation methods, exploring the peculiarities of each research group. Versatile and flexible features also allow studying cognitive and behavioral processes that are not observable with ordinary equipment. Finally, this custom equipment is also a low-cost alternatives to commercially available devices, making them a very interesting solution for cognition and behavioral research in animals.

## Supporting information

General Instructions

## ACKNOWLEDGMENTS

We would like to thank Grace Schenatto Pereira Moraes for helpful comments on the manuscript and Davi Barreto Mourão e Frederico José Barreto Mourão for scientific inspiration.

## AUTHOR CONTRIBUTIONS STATEMENT

Projected conceived and designed by P.A.A.J., F.A.G.M. and M.F.D.M.; P.A.A.J., F.A.G.M., M.C.L.A. and M.F.D.M. designed the hardware and code; P.A.A.J., performed sound and footshook validations; H.P.P., performed behavioral tests; L.O.G., performed behavioral analysis; F.A.G.M., H.P.P., performed surgery and electrophysiological records; F.A.G.M. and V.R.C. performed electrophysiological analysis. F.A.G.M. prepared figures; H.A.M supervised and contributed to refinements of the system; P.A.A.J., F.A.G.M. draft manuscript; M.F.D.M., H.P.P. and F.A.G.M. edited and revised manuscript; All authors approved the final version of manuscript.

