## Supplementary material for "A Custom Microcontrolled and Wireless-Operated Chamber for Auditory Fear Conditioning": General Instructions

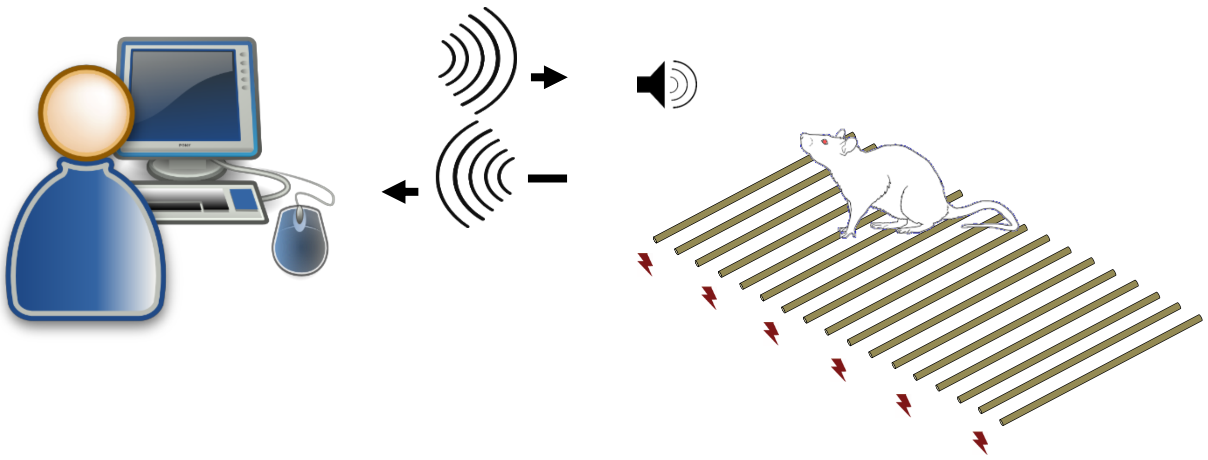

NNC

### Assembly Instructions

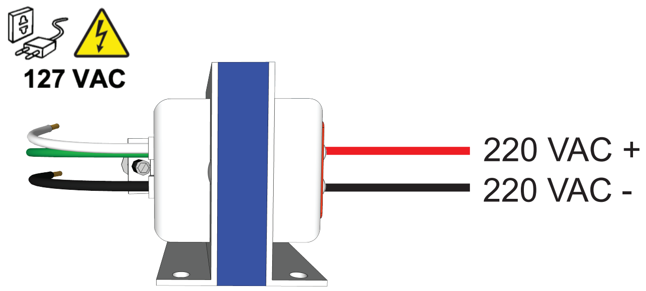


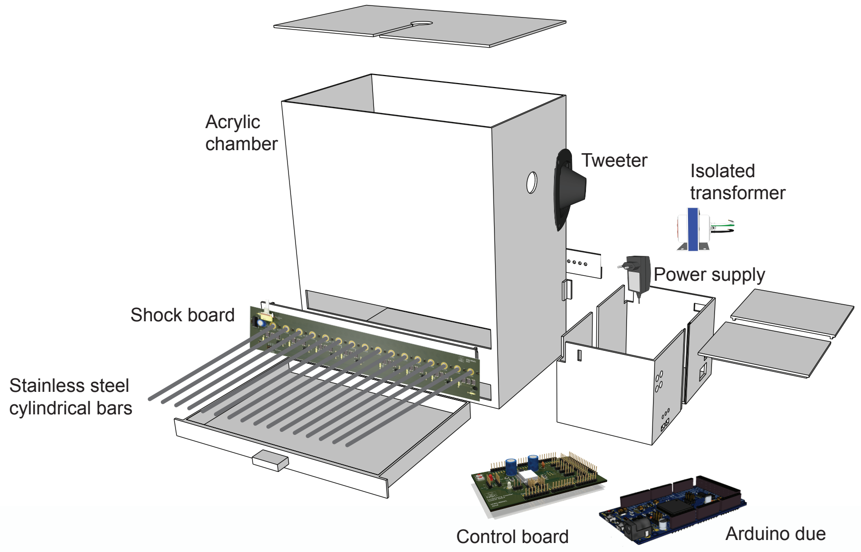


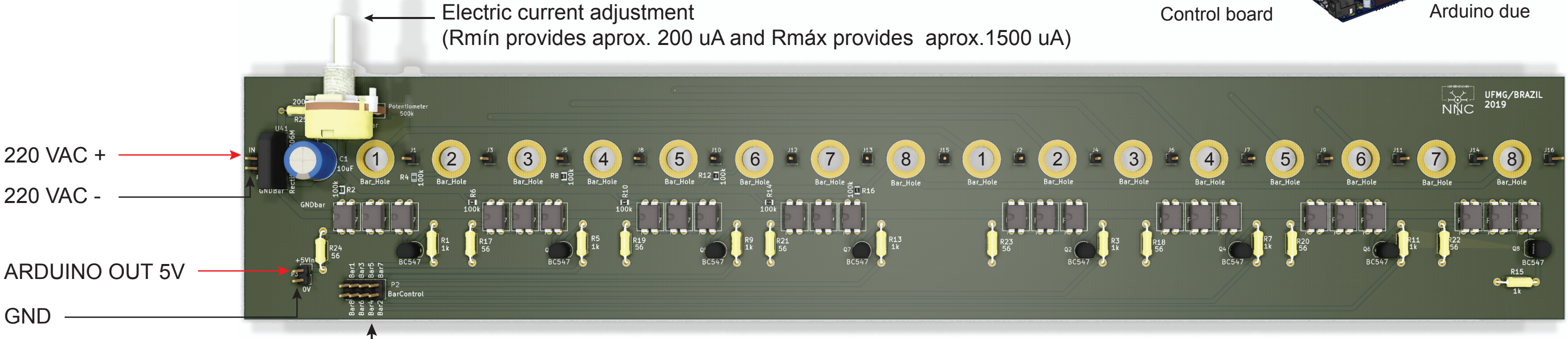


Buid in KiCad EDA 5.1.2 (http://www.kicad-pcb.org/)

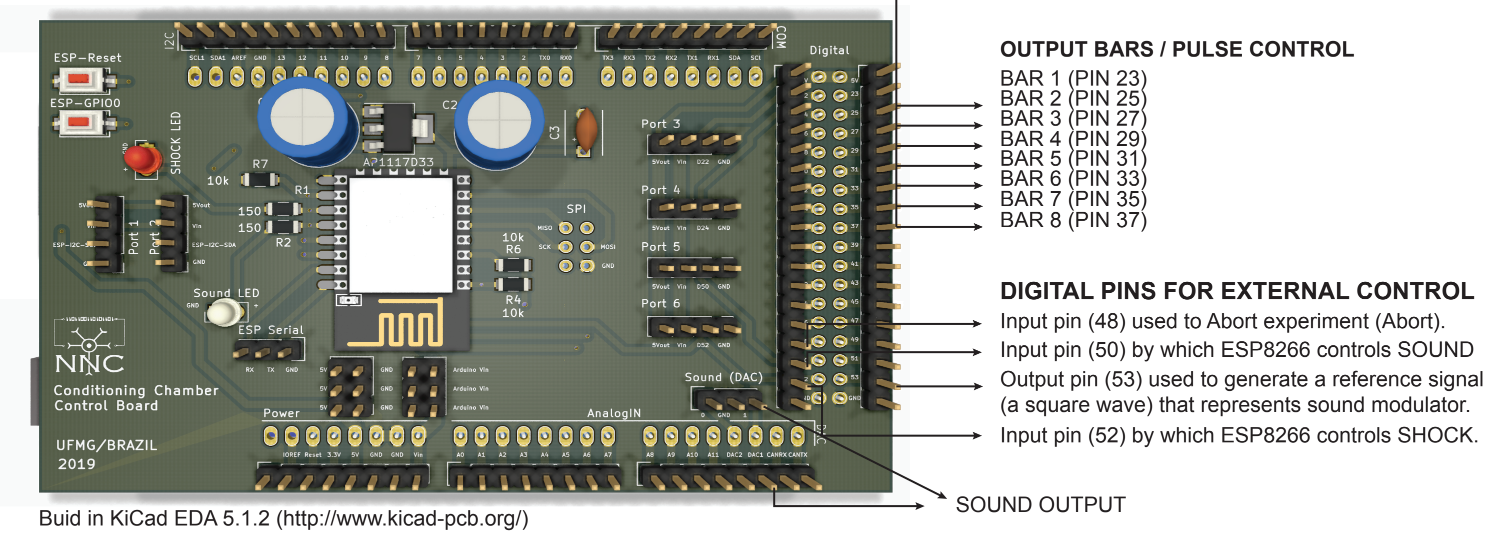


Buid in KiCad EDA 5.1.2 (http://www.kicad-pcb.org/)

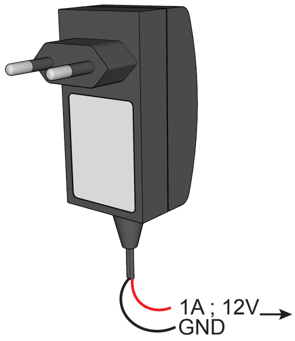


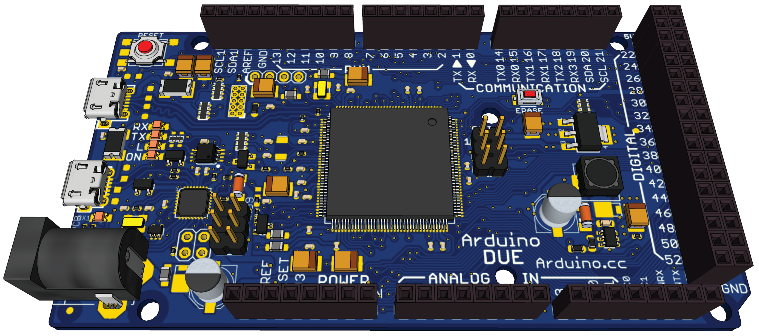


*https://store.arduino.cc/usa/due*
*https://3dwarehouse.sketchup.com/model/uef043628-8edb-4632-96bb-0f367bee29d8/Arduino-DUE?hl=en*

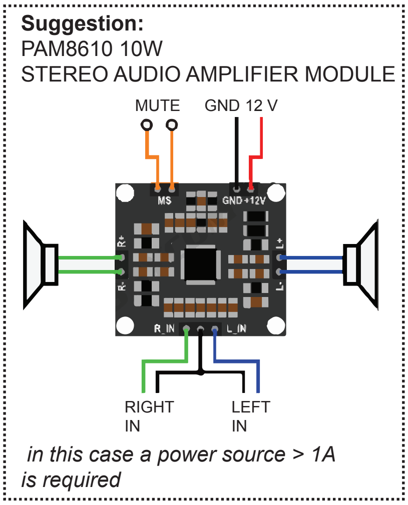


*https://hobbycomponents.com/audio/*
*664-pam8610-10w-stereo-audio-amplifier-module*

#### Original bar configuration for experiments with rats

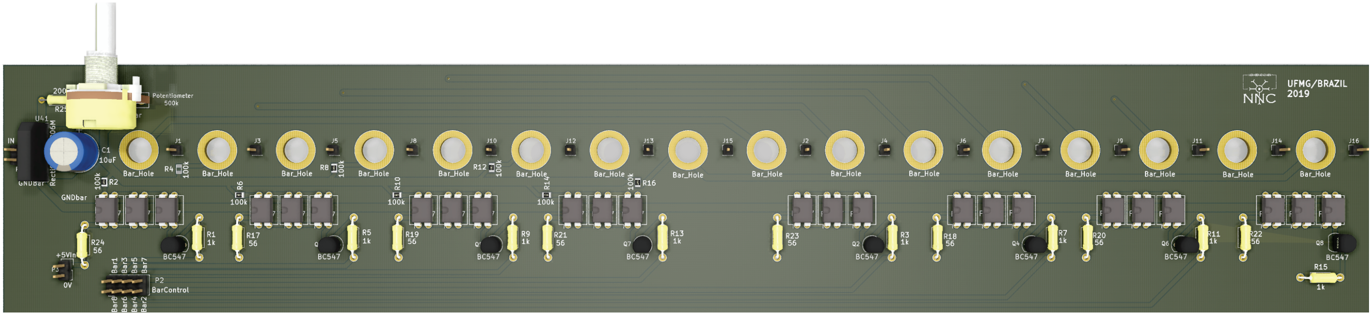

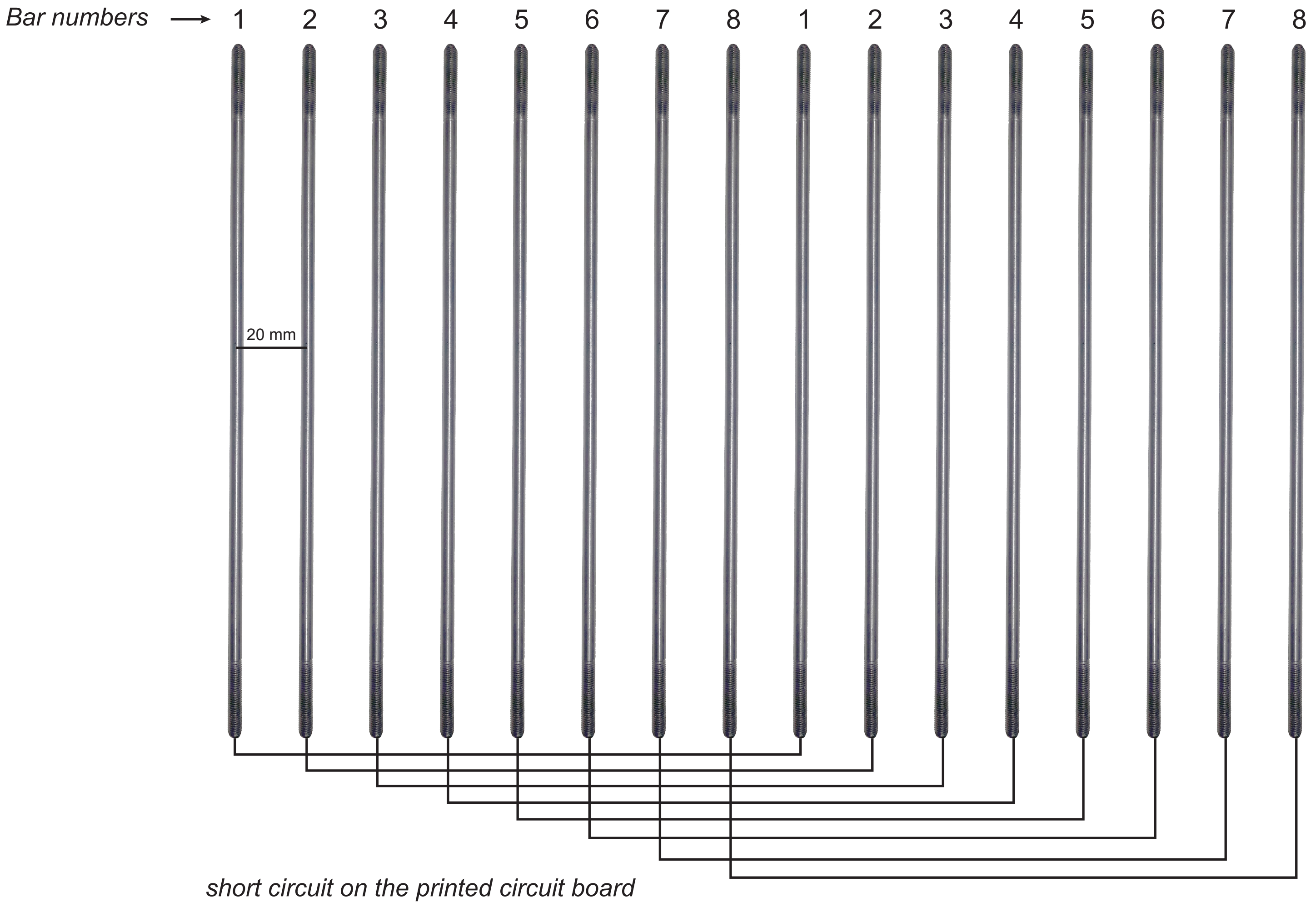

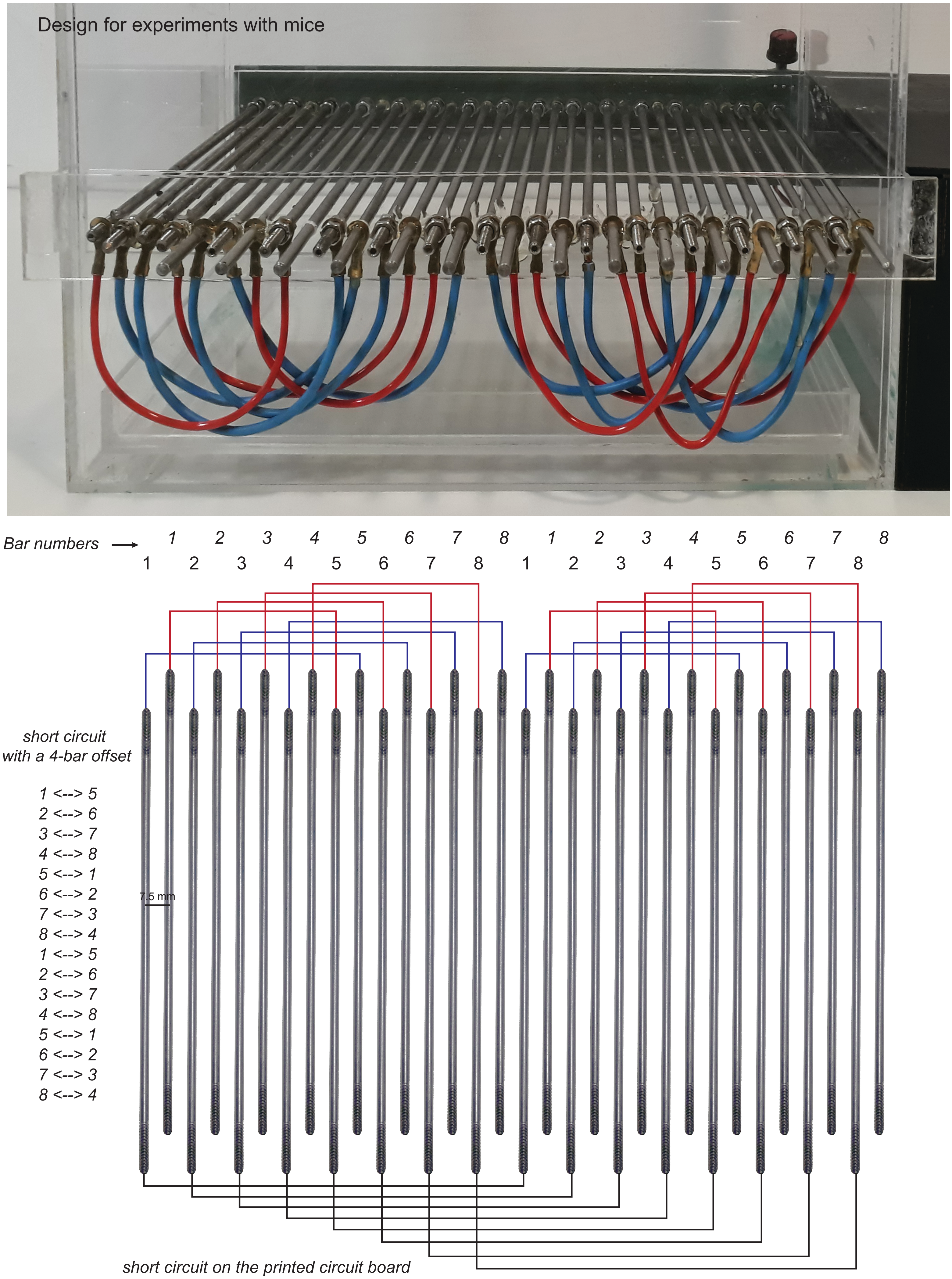


For experiments with mice the number of bars can be increased up to 32 and spaced 7.5 mm from each other. We suggest an external short circuit wired with a 4-bar offset. So that there are no repetitions in an 8-bar sequence covering the full animal extent.

**Power Supply**

| Component | Specification | Quantity |
|---|---|---|
| Isolated Transformer | 0.03KVA; $127V_{AC}$:$220V_{AC}$ | 1 |
| Power Supply | 1A; 12V | 1 |

**Two-layer printed circuit board with control circuit. It has ESP8266-12E footprint and an Arduino Due shield**

| Component | | Quantity |
|---|---|---|
| Arduino DUE | https://store.arduino.cc/usa/due | 1 |
| Module WI-FI ESP8266-12E | https://www.adafruit.com/product/2491 | 1 |
| FTDI Serial TTL-232 USB | https://www.adafruit.com/product/70 | 1 |

*TOP

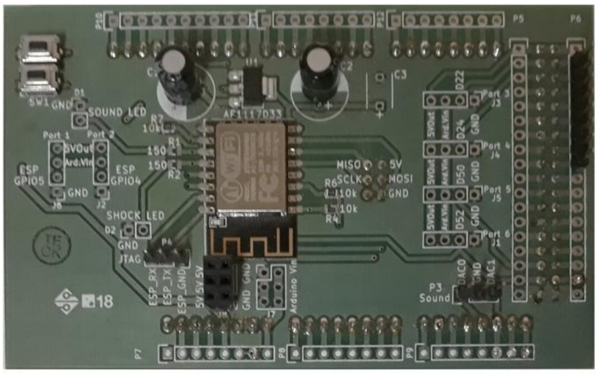

*BOTTOM

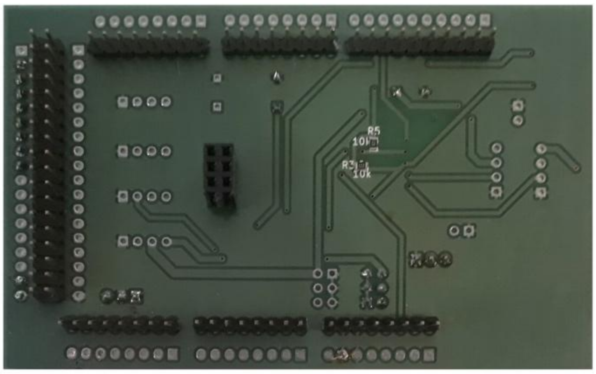

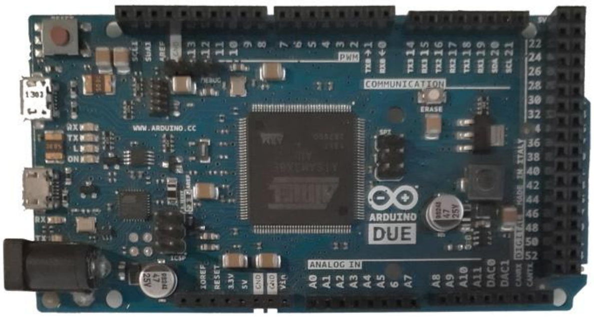


| Component | Model | Specification | Quantity |
|---|---|---|---|
| LDO Voltage Regulators | AP1117D33 | | 1 |
| Radial Electrolytic Capacitor | | 147uF, 550V | 2 |
| Resistor SMD | | 10kΩ 1% | 4 |
| Ceramic Capacitor | | 47uF, 50V | 1 |
| SMD Tactile Push button / Key Switch (2 x 6 x 2.5 mm) | KFC-A06 | | 2 |
| Pin Headers 2.54mm single (1 x 3 pins) | | | 8 |
| Pin Headers 2.54mm single (1 x 4 pins) | | | 6 |
| Pin Headers 2.54mm single (1 x 8 pins) | | | 5 |
| Pin Headers 2.54mm single (1 x 10 pins) | | | 1 |
| Dual Pin Headers 2.54mm single (2 x 18 pins) | | | 1 |

**Two-layer printed circuit board with power circuit. Attached to the chamber via 16 mounting holes, this board allows creating electric potential between the conductive bars.**

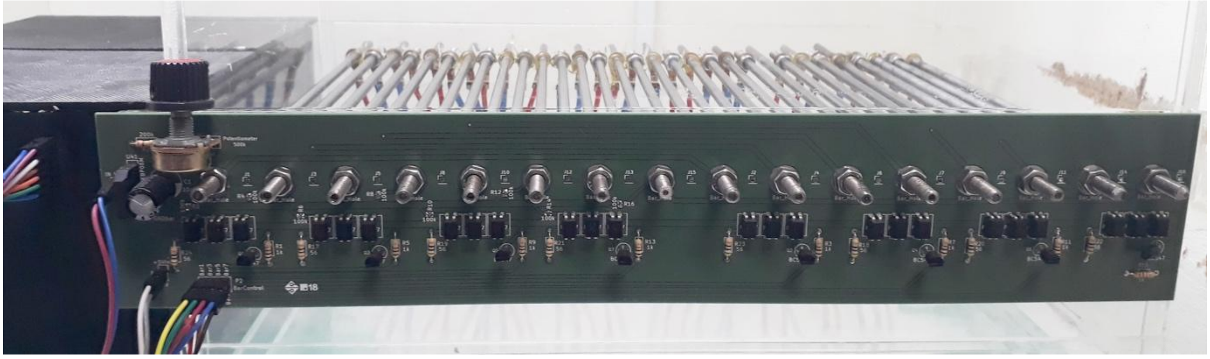

| Component | Model | Specification | Quantity |
|---|---|---|---|
| Resistor Through-hole | | 56Ω 5% 1/4W | 8 |
| Resistor Through-hole | | 1kΩ 5% 1/4W | 8 |
| Resistor Through-hole | | 200kΩ 5% 1/4W | 1 |
| Resistor SMD | | 100kΩ 1% 1/4W | 8 |
| Potentiometer -Type B Linear response curve | | 500kΩ | 1 |
| Radial Electrolytic Capacitor | | 10uF, 350V | 1 |
| Bridge Rectifier Diode | 2KBP06M | 2600V, 2A. | 1 |
| Optocouplers | PC817 4-pin | - | 24 |
| Transistors | BC547B | - | 8 |
| Pin Headers 2.54mm single (1 x 12 pins) | | | 1 |
| Dual Pin Headers 2.54mm single (2 x 4 pins) | | | 1 |
| Pin Headers 2.54mm single (1 x 2 pins) | | | 2 |
| Female Jumper wires | | | 12 |

**Stainless steel cylindrical bar and hex nuts**

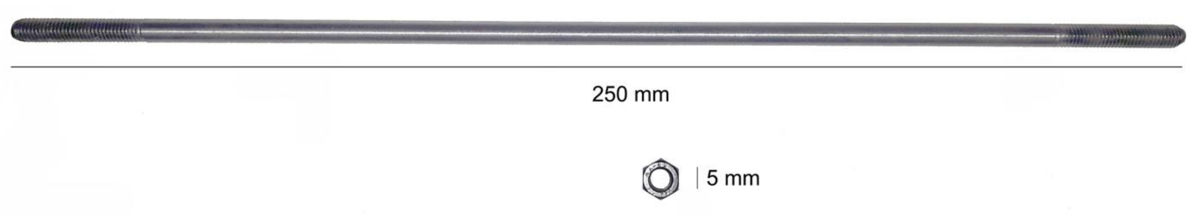


| Component | | Quantity |
|---|---|---|
| Stainless steel cylindrical bars | 250 x 5 mm | 32 (or 64) |
| Stainless steel hex nuts | 5 mm | 64 (or 128) |

#### Control Box

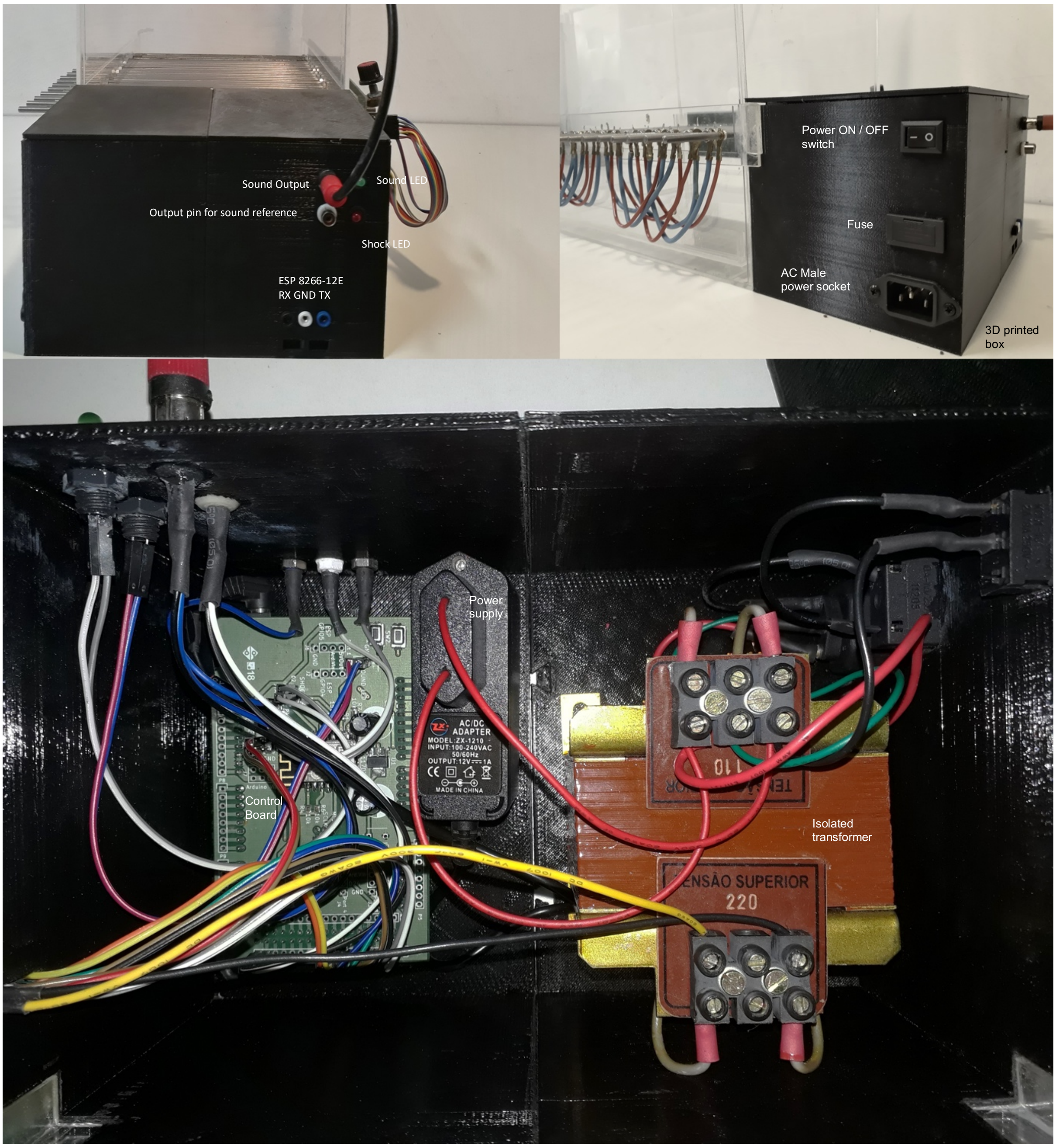


| Component | | Quantity |
|---|---|---|
| Power ON/OFF switch | 18.4 x 11.6 mm | 1 |
| AC Male computer power socket | Standard | 1 |
| Fuse Socket | - | 1 |
| Fuse 1A | - | 1 |
| Female RCA connector | - | 2 |
| Led | 5 mm | 2 |
| Female banana connector | 4 mm | 3 |
| Dual Pin Headers 2.54 mm single (2 x 6 pins) | | 1 |
| Female Jumper wires | | 12 |
